# In vitro resistance selection in *Acinetobacter baumannii* against the antimicrobial peptide TAT-RasGAP_317-326_ can cause cross-resistance to polymyxins through mutations in *pmrAB*

**DOI:** 10.1101/2025.01.22.634246

**Authors:** Emily Ritz, Tiffany Rossel, Nicolas Jacquier

## Abstract

Antibiotic resistance is a growing public health concern. In this context, there is an urgent need of alternative antimicrobial agents effective towards multidrug-resistant bacteria. Antimicrobial peptides (AMPs) are naturally occurring peptides, part of the first antimicrobial line of defence of a wide variety of organisms. AMPs were identified as a promising alternative to classical antibiotics with a low potential of resistance emergence towards them. However, increasing pieces of evidence indicate that resistance to AMPs may be more common than expected.

TAT-RasGAP_317-326_ is a semi-synthetic peptide first developed as an anticancer agent and later identified as an antibacterial agent. This peptide has a potent activity against Gram-negative bacteria, with a particularly low MIC (8 µg/ml) against *Acinetobacter baumannii*. While attempting to decipher the mode of action of TAT-RasGAP_317-326_, we performed in vitro resistance selection on *Escherichia coli* and only detected specific resistance to TAT-RasGAP_317-326_, without emergence of cross-resistance to other antimicrobial agents.

In this study, we performed a similar in vitro resistance selection assay using several *A. baumannii* strains. We repeatedly observed, upon selection with TAT-RasGAP_317-326_, the emergence of cross-resistance to polymyxins, a family of polypeptidic antibiotics used as last resort agents towards multiresistant bacteria. A majority of the cross-resistant strains we selected had mutations in the *pmrAB* operon. Importantly, some of these mutations were identical to mutations detected in polymyxins resistant clinical isolates. Such mutations are known to cause resistance to polymyxins through modifications of the charge and structure of lipopolysaccharides at the bacterial surface.

We thus show here that contact of *A. baumannii* with a semi-synthetic peptide structurally very different from polymyxins can induce the emergence of cross-resistance towards them. This indicates that caution should be taken with the clinical use of AMPs, since unexpected cross-resistance could emerge.

**Author Summary:** Antimicrobial peptides are currently deployed as an alternative treatment option towards multidrug resistant bacteria. However, little is known on the capacity of bacterial pathogens to develop resistance to these peptides. Here we show that the nosocomial pathogen *Acinetobacter baumannii* can develop resistance to the antimicrobial peptide TAT-RasGAP_317-326_ *in vitro*. In half of the cases, this was accompanied by cross-resistance to polymyxins, polypeptidic antibiotics used as last-resort treatment for multidrug resistant bacteria. We showed that this cross-resistance was generally caused by acquisition of mutations in the *pmrAB* genes, as observed in polymyxin-resistant clinical isolates. These mutations apparently cause changes in lipopolysaccharide structure, affecting the affinity of both TAT-RasGAP_317-326_ and polymyxins to the bacterial surface. Our results indicate that antimicrobial peptides should be used carefully, since they might induce cross-resistance to other antimicrobial agents. We observed this cross-resistance emergence with one peptide, TAT-RasGAP_317-326_ towards one pathogenic bacterium, *A. baumannii*. In the future, we need to determine whether such phenomena can happen with other bacterial species and other antimicrobial peptides.

## Introduction

Antibiotic resistance is a rising concern for modern medicine, rendering crucial the development of alternative treatment strategies. In this context, antimicrobial peptides (AMPs) have been proposed as such an alternative. AMPs are naturally occurring peptides produced by a wide variety of organisms as part of their first line of defence against pathogens (1). They possess very diverse sequences and sizes and can be produced either by ribosomes or via enzymatic complexes. These molecules can contain non-classical amino acids and be highly modified and even cyclized (2). Only a limited number of AMPs have been approved for clinical use (3). This is the case for polymyxins (colistin and polymyxin B), which are usually used as last-resort antimicrobial agents, despite potential important side effects (4). AMPs are proposed as promising antimicrobial agents since it is usually believed that the development of resistance towards them is very limited. This has been challenged by studies showing that bacteria can develop resistance to AMPs (5, 6). Resistance to AMPs seems generally specific to a subclass of AMPs and only limited cross-resistance could be observed (7, 8). However, it was recently shown that resistance to colistin found in clinical strains can cause moderate cross-resistance to host AMPs, inducing an increased virulence of the resistant strains (9). Nevertheless, multidrug resistant (MDR) bacteria are usually sensitive to AMPs (10), indicating that broad cross-resistance to both classical antibiotics and AMPs is unlikely. Cross-resistance development was mainly studied in the model organism *Escherichia coli* (8). Whether these observations can be extended to other pathogenic bacteria is still an open question.

*Acinetobacter baumannii* is a Gram-negative opportunistic human pathogen. It is a common agent of nosocomial infections that colonizes both the respiratory and urinary tract. A strong advantage of *A. baumannii* is its capacity to form biofilms on biotic and abiotic surfaces (11). These biofilms can be highly tolerant to antimicrobial agents and disinfectants. *A. baumannii* shows high genome plasticity that allows this bacterium to rapidly gain resistance to antibiotics such as carbapenem (12). For this reason, carbapenem-resistant *A. baumannii* was highlighted by the WHO as a critical priority pathogen for research and development of new antibiotics. Polymyxins are used as last-resort antibiotics against these carbapenem-resistant *A. baumannii*. Nevertheless, emergence of polymyxin-resistant strains of *A. baumannii* has also been documented (13). Resistance to polymyxins usually involves mutations in genes encoding the two-component system PmrAB, which induce modifications of the lipopolysaccharide (LPS), also known as lipooligosaccharide (LOS) in *A. baumannii*, via the addition of positive charges to Lipid A (14). AMPs have been pointed as promising antimicrobial agents towards *A. baumannii* (15). However, little is known on the potential of *A. baumannii* to acquire resistance towards these AMPs.

TAT-RasGAP_317-326_ was first developed as an anticancer peptide and was later characterized as an antibacterial agent (16). It showed promising activity towards Gram-negative bacteria, especially towards both *A. baumannii* ATCC19606 strain (MIC of 8 µg/ml) and MDR clinical isolates (MIC of 16 µg/ml) (16). This peptide also had inhibitory activity against an *in vitro* model of *A. baumannii* biofilm, with a minimal biofilm inhibition concentration between 32 µg/ml and 64 µg/ml when used alone, and 4 µg/ml when used in combination with 4 µg/ml of meropenem (17, 18). It also partially affects the viability of mature biofilm alone or in combination with meropenem (17, 18). In addition, efficiency of TAT-RasGAP_317-326_ towards *A. baumannii* was further increased by combining it with a selection of antibiotics. Combination with gentamicin or meropenem resulted in synergism towards planktonic bacteria and increased efficiency towards biofilm (17).

In this study, we investigate the capacity of different *A. baumannii* strains to acquire resistance to TAT-RasGAP_317-326_ *in vitro* and whether acquisition of resistance to this peptide caused the appearance of cross-resistance to other antimicrobial agents. We show that cross-resistance to polymyxins emerges without a strong impact on bacterial fitness and virulence. Cross-resistance is caused by acquisition of mutations in the genes encoding the PmrAB two-component system. Some of the mutations we detected are identical to documented mutations found in clinical strains of *A. baumannii* resistant to polymyxins. These results indicate that treatment with an AMP, which structure is totally different from polymyxins could still cause the emergence of resistance to polymyxins in *A. baumannii* and highlights the need for caution regarding the clinical application of AMPs.

## Results

### *In vitro* selection of resistance towards TAT-RasGAP_317-326_ in *A. baumannii* recurrently results in cross-resistance to polymyxins

In order to study the potential of *A. baumannii* isolates to develop resistance to TAT-RasGAP_317-326_, we selected 9 isolates with diverse backgrounds and antibiotic resistance profiles, comprising two well studied ATCC strains and seven clinical isolates (Table 1). These strains were subjected to eight passages in liquid culture with increasing concentration of the antimicrobial peptide TAT-RasGAP_317-326_, starting with subinhibitory concentrations and then increasing concentrations when growth was observed, similarly as was already performed with other bacterial species (19, 20). In parallel, the same experiment was performed on the ATCC 19606 strain using polymyxin B and tetracycline, respectively. Resistance profiles of the selected strains were then measured (Figure 1 and Tables 2 and 3). A limited but consistent increase in MIC of TAT-RasGAP_317-326_ was observed in a large majority of strains selected with this peptide (Fig. 1A). Interestingly, development of resistance to this peptide was linked, in some cases to the development of cross-resistance to polymyxins (polymyxin B and colistin), but not to the AMP melittin or to antibiotics gentamicin or tetracycline (Table 2, resistance development highlighted in bold). In contrast, incubation of ATCC 19606 strain with increasing concentrations of polymyxin B or tetracycline induced the emergence of resistance specific to polymyxins or tetracycline, respectively (Figure 1B-C and Table 3). We did not observe appearance of any potent collateral sensitivity, as was observed in other studies (7).

**Table 1:** List of *A. baumannii* strains used in this study. Minimal inhibitory concentrations (MICs) of TAT-RasGAP_317-326_ (TAT), polymyxin B (PolB), colistin (Col), melittin (Mel), gentamicin (Genta) and tetracycline (Tetra) against the indicated strains were measured in duplicates. Both values are indicated when different results were obtained.

| Isolate name | MIC TAT | MIC PoIB | MIC Col | MIC Mel | MIC Genta | MIC Tetra | Ref. |
| --- | --- | --- | --- | --- | --- | --- | --- |
| ATCC19606 | 8-16 | 1-2 | 2 | 16 | 32 | 1 | ATCC |
| ATCC17978 | 8 | 2-4 | 0.5-1 | 8 | 4 | 0.5 | ATCC |
| Ab1 | 8-16 | 1-2 | 4 | 8-16 | >256 | >256 | Heulot |
| Ab2 | 16 | 2-4 | 0.25 | 8-16 | >256 | 1 | Heulot |
| Ab3 | 16 | 1-2 | 2-4 | 32 | >256 | >256 | Heulot |
| Ab31 | 16 | 2 | 4-8 | 16-64 | 16 | 1 | Leshkasheli |
| Ab33 | 16 | 2-4 | 4-8 | 32 | >256 | >256 | Leshkasheli |
| Ab44 | 16 | 1 | 2 | 8-16 | 64 | 2 | Leshkasheli |
| Ab73 | 8-16 | 1-2 | 1 | 8-16 | 8 | 0.5-1 | Leshkasheli |

**Table 2:** MICs of strains selected with TAT-RasGAP_317-326_. Minimal inhibitory concentrations (MICs) of TAT-RasGAP_317-326_ (TAT), polymyxin B (PolB), colistin (Col), melittin (Mel), gentamicin (Genta) and tetracycline (Tetra) against the indicated strains were measured in duplicates. Both values are indicated when different results were obtained. Bold values indicate that the measured MICs are at least 3 times higher than the parental strain.

| Isolate name | MIC TAT | MIC PoIB | MIC Col | MIC Mel | MIC Genta | MIC Tetra |
| --- | --- | --- | --- | --- | --- | --- |
| ATCC19606 | **48** | 4-6 | 1-2 | 8-16 | 64 | 0.5 |
| ATCC17978 | **64** | **64** | **>128** | 16 | 4-8 | 0.5 |
| Ab1 | **32-48** | 4-6 | 4-8 | 16-32 | >256 | 128-256 |
| Ab2 | **48** | **32** | **128** | 8-16 | >256 | 2 |
| Ab3 | **64** | 3-4 | 2-4 | 32-64 | >256 | 256 |
| Ab31 | 24-32 | 4 | 2-8 | 32 | 8 | 0.5 |
| Ab33 | **48** | **32** | **>128** | 16 | >256 | >256 |
| Ab44 | **64** | **32-64** | **>128** | 32 | 64 | 2-4 |
| Ab73 | **48-64** | **8** | **128** | 16 | 8 | 0.5-1 |

**Table 3:** MICs on the strains selected with tetracycline and polymyxin B. Minimal inhibitory concentrations (MICs) of TAT-RasGAP_317-326_ (TAT), polymyxin B (PolB), colistin (Col), melittin (Mel), gentamicin (Genta) and tetracycline (Tetra) against the indicated strains were measured in duplicates. Both values are indicated when different results were obtained. Bold values indicate that the measured MICs are at least 3 times higher than the parental strain.

| Isolate name | MIC TAT | MIC PoIB | MIC Col | MIC Mel | MIC Genta | MIC Tetra |
| --- | --- | --- | --- | --- | --- | --- |
| ATCC19606 PoIB | 24 | **128** | **>128** | 8 | 64-128 | 0.5-1 |
| ATCC19606 Tetra | 32 | 2 | 2 | 4-16 | 32 | **64** |

**Figure 1.**
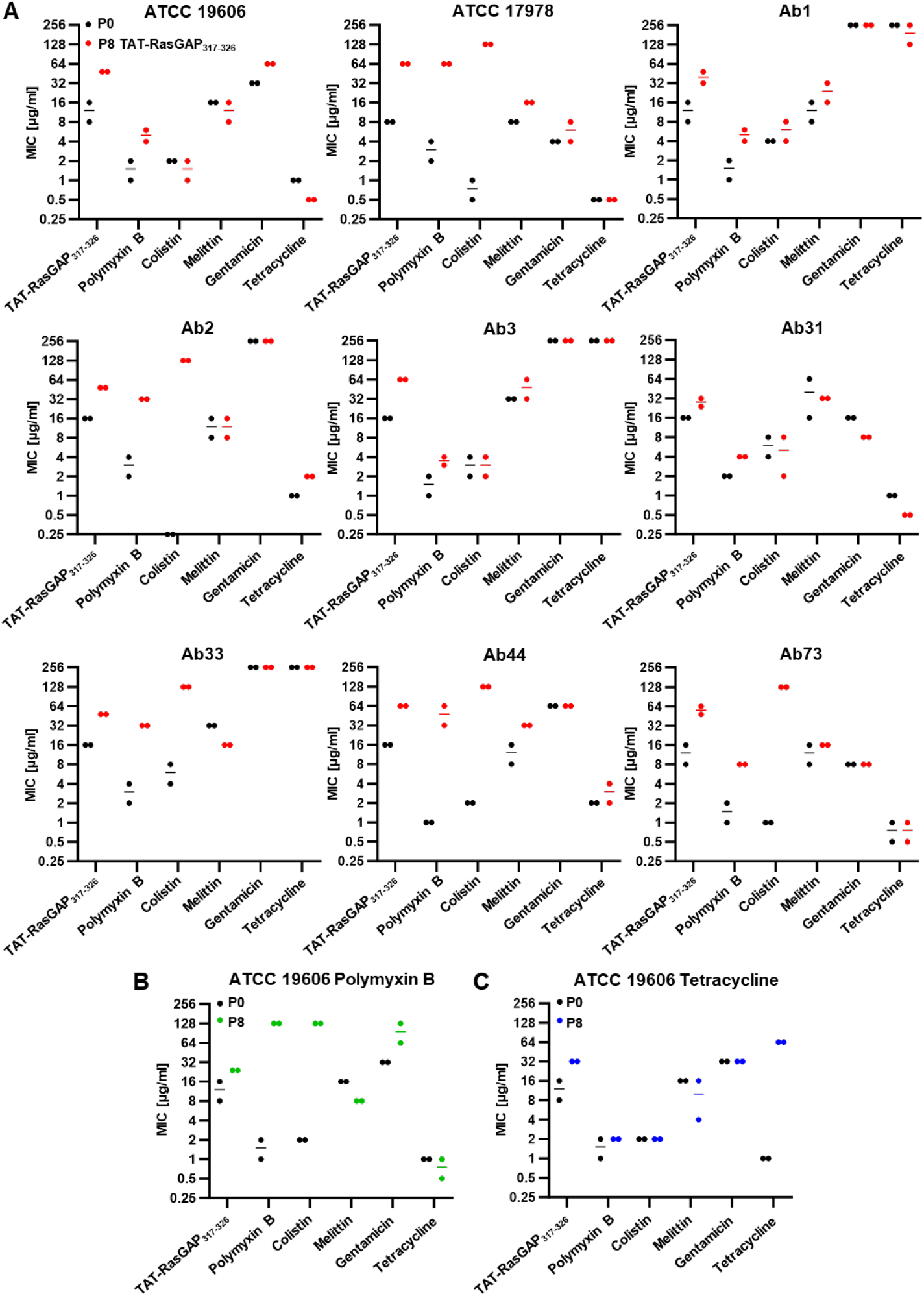
*In vitro* selection resistance in the presence of TAT-RasGAP_317-326_ can induce both specific resistance and cross-resistance to polymyxins in different strains of *Acinetobacter baumannii*. The indicated strains were incubated with increasing concentrations of TAT-RasGAP_317-326_ **(A)**, polymyxin B **(B)** or tetracycline **(C)** for a total of 8 passages to obtain resistant isolate (P8). Minimal Inhibitory Concentrations of the indicated antimicrobial agents were determined for both P8 and parental strains (P0). Exact values are shown in Table 1 (P0) and Table 2 (P8 with TAT-RasGAP_317-326_) and Table 3 (P8 selected with polymyxin B or tetracycline). Measurements were performed in duplicates, and lines represent the average of the two values, when different.

### Development of resistance towards TAT-RasGAP_317-326_ does not cause collateral fitness defects

Acquisition of resistance is sometimes linked to a fitness cost. We thus measured growth rate of resistant mutants and compared it with growth of the parental strains. We did not observe any growth defect, both in LB (Figure 2) and in LB supplemented with NaCl or glucose (Figures S1 and S2). Moreover, we did not observe any important morphological changes of planktonic bacteria between the resistant isolates and their parental counterparts (Figure S3).

**Figure 2.**
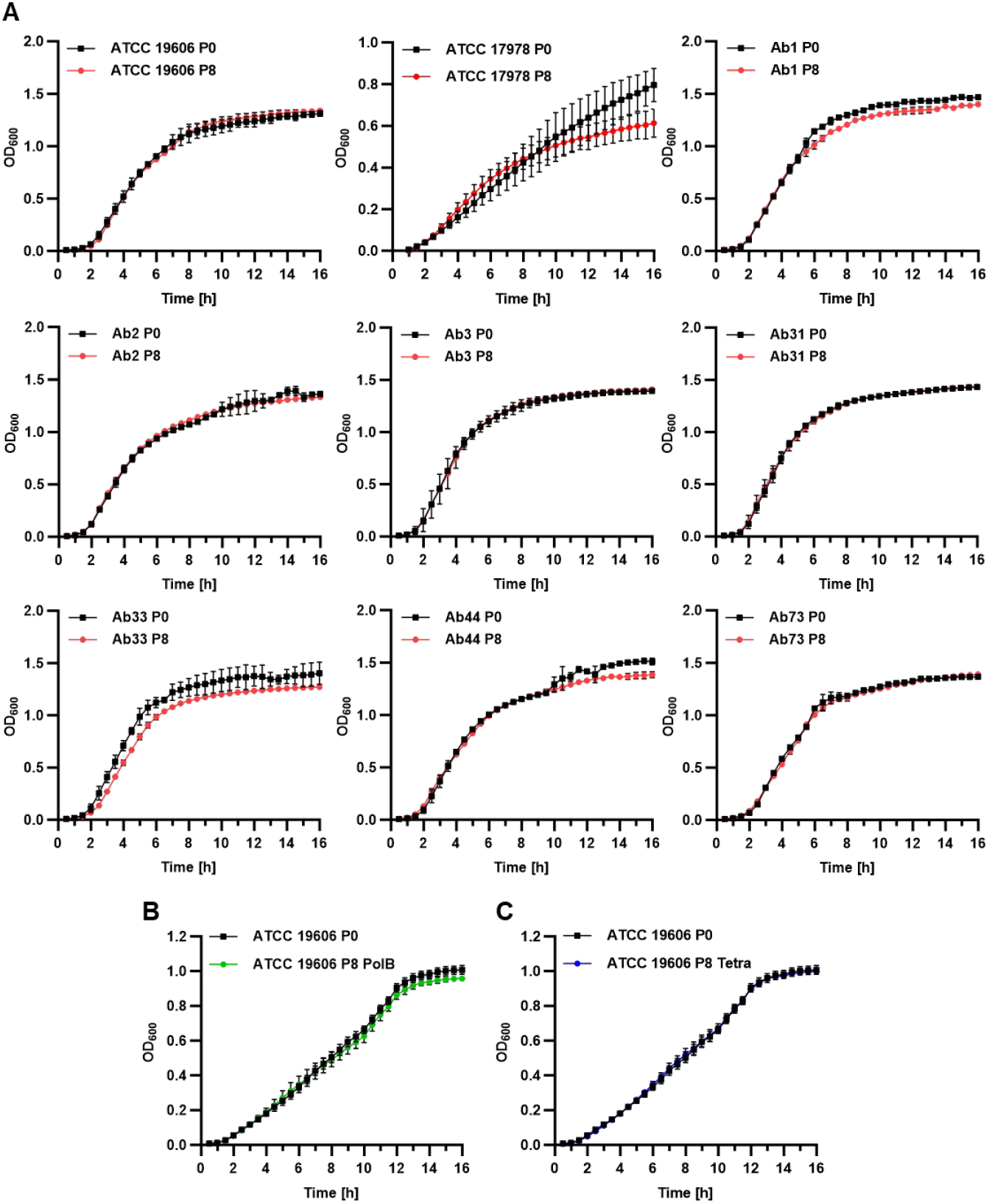
Resistance acquisition does not induce *in vitro* growth defects. P0 and P8 *A. baumannii* isolates upon selection with TAT-RasGAP_317-326_ **(A)**, polymyxin B **(B)** or tetracycline **(C)** were grown overnight in LB and diluted to an OD_600_ of 0.01 in fresh medium. Growth was then assessed by OD_600_ measurement each 30 minutes for a total of 16 hours. Values are the average of three experiments. Error bars represent standard deviation of the three replicates.

Since *A. baumannii* is well known to form biofilm, we decided to test whether development of resistance affects the capability of the different strains to form biofilm, using an *in vitro* model we already used in the past (17). Using crystal violet staining, we quantified the amount of biofilm formed by the different resistant strains and compared them with their parental counterparts (Figure 3A). We did not observe clear trends when comparing resistant and sensitive strains. In a further step, we formed biofilm on glass coverslips, labelled them with live-dead staining and performed confocal microscopy (Z-stacks) in order to observe the morphology and thickness of the different biofilms (Figure 3B and Figure S4). Again, no strong differences in biofilm formation capacities were observed, indicating that acquisition of resistance did not induce defects in biofilm formation in the *A. baumannii* strains we investigated. Taken together, these results indicate that acquisition of both resistance specific to TAT-RasGAP_317-326_ and cross-resistance to polymyxins does not strongly influence bacterial fitness and morphology.

**Figure 3.**
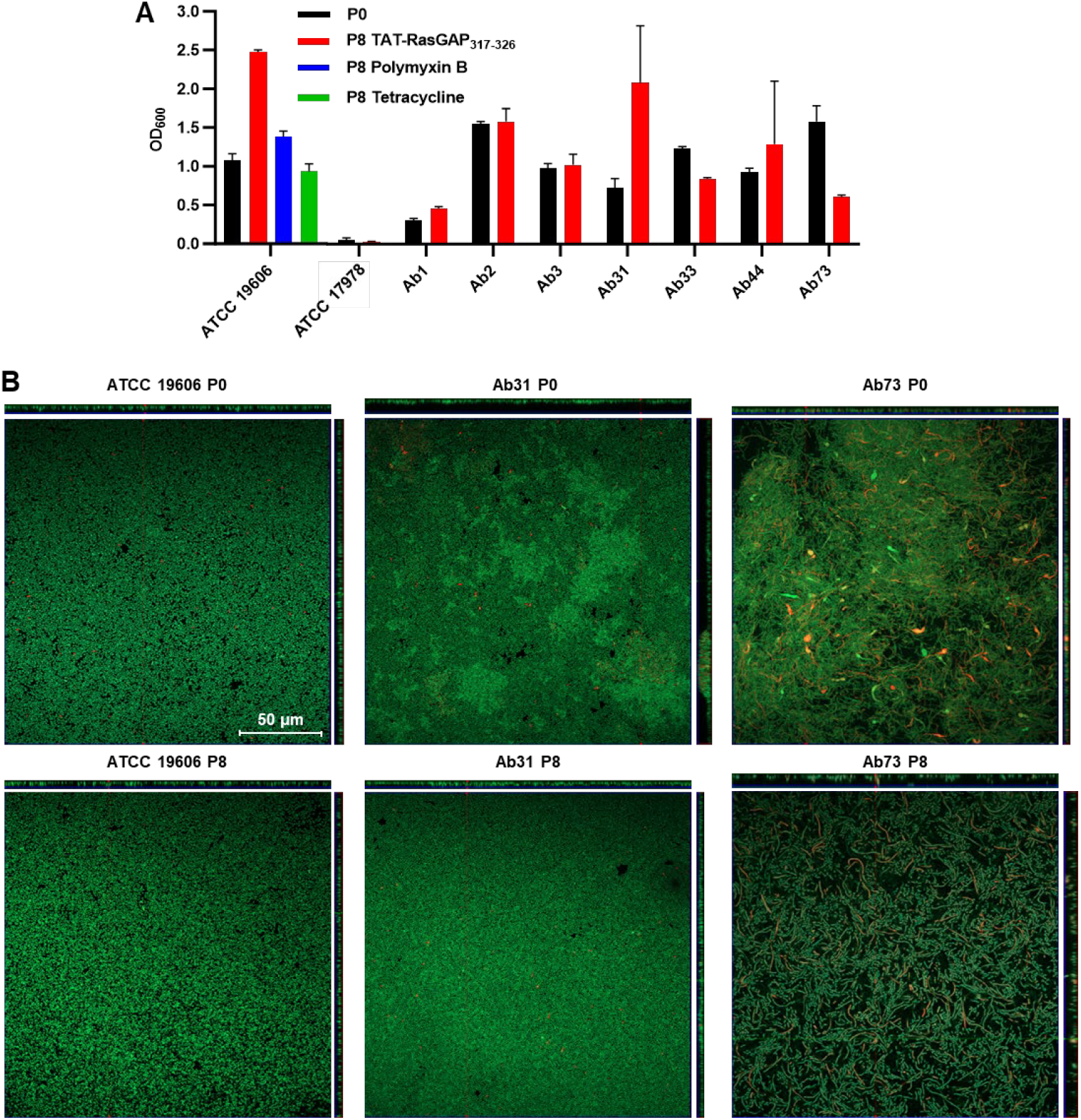
Biofilm formation and virulence are not strongly affected in *A. baumannii* isolates resistant to TAT-RasGAP_317-326_. **(A)** *In vitro* biofilm formation was assessed for both P0 and P8 isolates, by quantifiying biofilm biomass through crystal violet staining. **(B)** The effect of resistance acquisition on the structure of the biofilm was assessed by live-dead staining followed by Z-stacks acquisition with a confocal microscope. Reconstruction of Z projections are shown on the top and on the right for each strain. Three representative strains are shown.

### Emergence of cross-resistance does not depend on a specific genetic background

We observed that cross-resistance to polymyxins emerged only in 5 of the 9 strains we tested. Since the resistance selection was performed only once per strain, it was impossible at that stage to determine whether cross-resistance can emerge only in a subset of strains. We thus repeated resistance selection for all the strains in triplicate. We measured the MICs of TAT-RasGAP_317-326_ and polymyxin B on the resulting strains (Table 4 and Figure S5). We could detect robust increase of MIC of TAT-RasGAP_317-326_ (more than three-fold increase) upon selection in 13 of the 27 selected isolates (Table 4, highlighted in bold). In contrast, MIC of TAT-RasGAP_317-326_ towards some replicates was not very different compared to the parental strain, indicating that efficiency of resistance emergence can vary. Selection of resistance to TAT-RasGAP_317-326_ was again linked, in 16 of the 27 isolates, to cross-resistance towards polymyxin B. This appears to occur stochastically, independently of the strain genetic background, since cross-resistance was detected at least once in all strains apart from Ab31. It should be noted that in both experiments, we were not able to obtain a robust MIC increase in the Ab31 strain, indicating that this strain may have lower capabilities to become resistant to TAT-RasGAP_317-326_. Furthermore, in order to determine whether resistance acquisition was stable, we passaged ATCC19606 resistant isolates for a total of 8 times in the absence of any antimicrobial agent. Resistance to polymyxins was unchanged after these 8 passages, indicating that resistance could not be lost (Figure S6).

**Table 4:** MICs of strains selected with TAT-RasGAP_317-326_. Minimal inhibitory concentrations (MICs) of TAT-RasGAP_317-326_ (TAT) and polymyxin B (PolB) against the indicated strains were measured in duplicates. Both values are indicated when different results were obtained. Values in bold indicate that the measured MICs are at least 3 times higher than the parental strain.

| Isolate name | MIC TAT |  |  | MIC PolB |  |  |
| --- | --- | --- | --- | --- | --- | --- |
| Replicate | b | c | d | b | c | d |
| ATCC19606 | 16-64 | **32-64** | **32-64** | 4-6 | **64** | 2-4 |
| ATCC17978 | 16-32 | **64-96** | **48-64** | 0.5-1 | **32-64** | 0.5 |
| Ab1 | **128->128** | **64** | 32 | **128->128** | **>128** | 4 |
| Ab2 | **32-64** | 16-64 | **64** | 1-2 | **64** | **64** |
| Ab3 | 16-32 | 32 | **64** | 4-8 | **32** | **128->128** |
| Ab31 | 24 | 24-32 | 16-32 | 2-4 | **64-128** | 4-8 |
| Ab33 | **48** | **128** | **48** | **>128** | **128->128** | **>128** |
| Ab44 | 24-32 | 32 | **64** | **16** | **32** | **128->128** |
| Ab73 | 12 | 12-16 | 16 | 4 | 2 | **32-64** |

### Cross-resistance emergence is generally linked to mutations in *pmrAB*

In order to better understand the mechanisms involved in the acquisition of cross-resistance, we performed whole genome sequencing on the resistant strains obtained in the first experiment (Table 2) and compared their genome with the corresponding parental strains. Mutations that were identified and that result in amino acid changes are shown in Table S1. Some of the detected mutations in isolates specifically resistant to TAT-RasGAP_317-326_ affect outer membrane proteins (OmpA, BauA), while others might influence gene expression regulation (TopA). Strain Ab1 is potentially a hypermutator strain since it acquired up to 20 mutations, while others only acquired between 1 and 4 mutations (Table S1).

Interestingly, four isolates out of the five that developed cross-resistance to polymyxin B acquired a mutation in the *pmrAB* operon. This operon encodes the two-component system PmrAB, which regulates LOS modifications, in particular addition of the cationic sugar L-4-aminoarabinose to the lipid A phosphate group of LOS (14). Mutations in PmrAB were already identified in polymyxin-resistant clinical isolates (21).

To further confirm the implication of the PmrAB two-component system in cross-resistance, we amplified *pmrA* and *pmrB* by PCR from all the resistant strains obtained and performed Sanger sequencing to identify potential mutations in these two genes. We could thus find mutations in *pmrA* or *pmrB* in 18 of the 21 cross-resistant isolates. In comparison, none of the specific resistant isolates had any mutations in these two genes (Figure 4A). We then mapped the mutations we detected on PmrA and PmrB protein domains and searched the literature to verify whether some mutations were already described. Indeed, 7 out of the 11 mutated sites we detected have already been found in clinical *A. baumannii* isolates resistant to colistin (21) (Figure 4B, mutations already described in literature are highlighted in bold).

**Figure 4.**
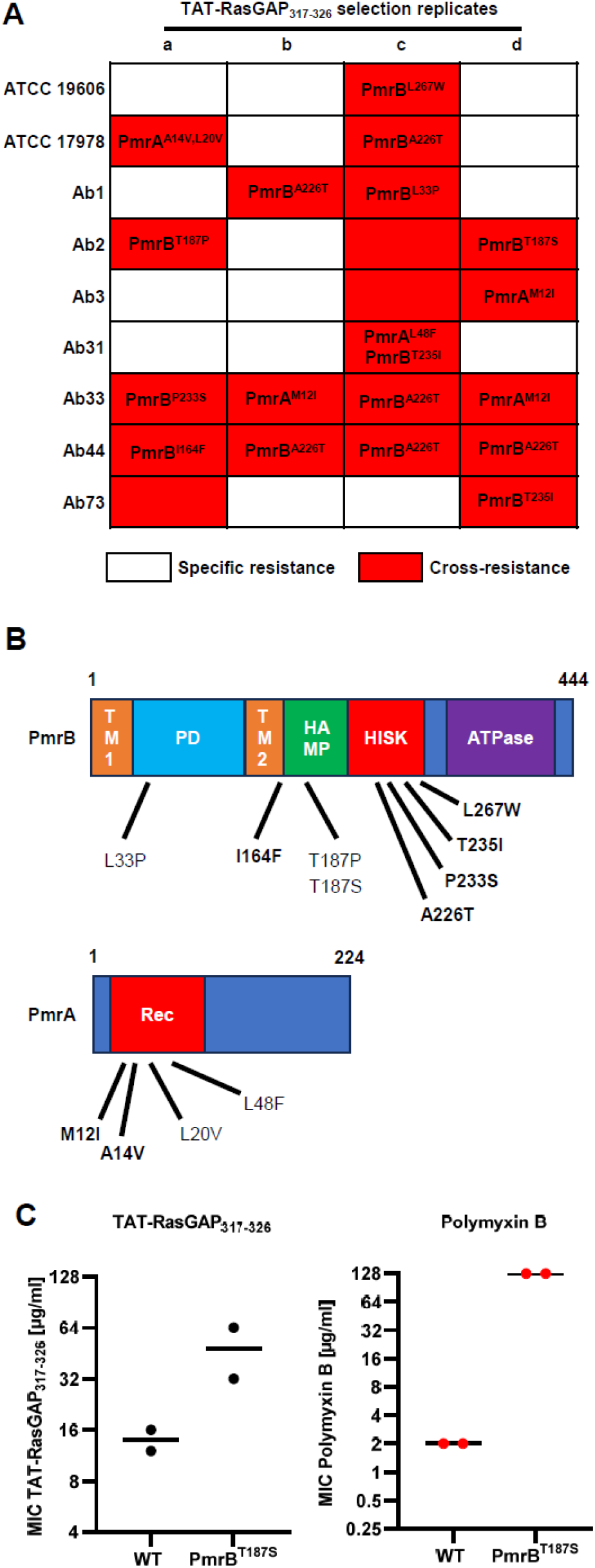
Emergence of cross-resistance to polymyxins upon selection with TAT-RasGAP_317-326_ mostly occurs through point mutations in *pmrAB*. **(A)** Resistant isolates depicted in Table 2 and Table 4 are classified as bearing a specific resistance (white) or cross-resistance (red). Sequencing of *prmA* and *pmrB* genes was performed on all strains and mutations found upon comparison with the parental strains are shown. **(B)** Amino acid changes caused by mutations found in cross-resistant strains were mapped on the protein domains of PmrA and PmrB. Mutations already described in the literature are highlighted in bold. **(C)** The PmrB_T187S_ mutation is sufficient to cause cross-resistance to TAT-RasGAP_317-326_ and polymyxin B. Targeted mutagenesis using a CRISPR-Cas9 based method was performed in an ATCC17978 wild-type strain. MICs of the indicated antimicrobial agents towards the mutant and the wild-type strain were measured in duplicates. Black lines represent the average of the replicates.

### Mutation in *pmrB* is sufficient to cause cross-resistance to TAT-RasGAP_317-326_ and polymyxin B

In order to confirm that such mutations are indeed the cause of the cross-resistance to TAT-RasGAP_317-326_ and polymyxin B, we used a CRISPR-Cas9 based method to insert a single mutation in a wild-type background (22). Using this method, we introduced the T187S mutation in the ATCC 17978 strain, which was sufficient to cause an increase in MIC of both TAT-RasGAP_317-326_ and polymyxin B towards this strain (Figure 4C).

### Cross-resistance sometimes appears without mutations in *pmrAB*

In three cross-resistant isolates, the phenotype could not be attributed to mutations in *pmrAB* (Ab2c, Ab3c, Ab73a). We already had the genome sequence of Ab73a (Table S1). We thus performed whole genome sequencing of the two isolates Ab2c and Ab3c and identified mutations by comparing with their respective parental strains (Table S2).

Four mutations could be detected in strain Ab73a. Two of them caused a change of one amino acid in the proteins BamA and LptC, respectively. A third caused the appearance of a premature stop codon, producing a truncated form of PlcD, and the last one caused a frameshift in the gene encoding PldA. We recently identified a potential role for the essential outer membrane insertase BamA as a receptor for TAT-RasGAP_317-326_ in *E. coli* (23). LptC is a periplasmic protein of the LOS transporter machinery. PlcD and PldA are phospholipases. Interestingly, inhibition of phospholipase function was shown to improve fitness of a LOS-deficient mutant in *A. baumannii* (24). It is thus possible that inactivation of phospholipases in the Ab73a strain compensates some defect in LOS transport caused by the LptC mutation.

A total of 10 mutations was detected in Ab2c strain, mainly affecting uncharacterized proteins. However, point mutations were detected to cause amino acid changes in UvrA, a thiamine permease and BasI, an acinetobactin biosynthetic protein. Finally, the start codon of the *crp* gene, encoding the cyclic AMP receptor protein, was mutated. This is expected to induce a depletion of the Crp protein. Crp is an important transcriptional regulator (regulating over 180 genes in *E. coli* (25)), which is regulated by cyclic-AMP binding. Cyclic-AMP has recently been identified as an important global virulence regulator in *A. baumannii* (26). Depletion of Crp might thus strongly affect cellular properties and thus lead to cross-resistance phenotype.

Concerning the Ab3c strain, only three single point mutations were detected in two genes, encoding uncharacterized proteins: a AAA family ATPase and an integrase family protein. Because these two proteins are uncharacterized, we cannot explain their role in cross-resistance. Overall, PmrAB-independent cross-resistance phenomena need to be further investigated in the future.

### Emergence of cross-resistance does not affect bacterial virulence

Since we observed no consistent influence of cross-resistance development on bacterial fitness *in vitro*, we decided to take advantage of the specific and cross-resistant isolates we obtained in the same background to test their virulence in an *in vivo Galleria mellonella* model. We decided to limit our study to the well described ATCC 19606 and ATCC 17978 strains. Indeed, infection models of *G. mellonella* using these two strains have been described in the literature (27, 28). We thus infected groups of *G. mellonella* larve with equivalent quantities of A. baumannii (10^6^ bacteria per larva), either sensitive (P0) specifically resistant to TAT-RasGAP_317-326_ (ATCC19606 P8a and ATCC 17978 P8b) or cross-resistant to TAT-RasGAP_317-326_ and polymyxins (ATCC 19606 P8c and ATCC 17978 P8a). Injection of all strains caused a significant drop in *G. mellonella* viability in comparison to the PBS injected control (Figure 5). No significant difference was observed between the different ATCC 19606 isolates (Figure 5A). A limited but significant increase of viability was observed for ATCC 17978 P8b strain, which shows specific resistance to TAT-RasGAP_317-326_ (Figure 5B). Taken together, these results do not show a significant effect of acquisition of cross-resistance on *A. baumannii* virulence. However, virulence might be lowered in some cases by acquisition of specific resistance to TAT-RasGAP_317-326_.

**Figure 5.**
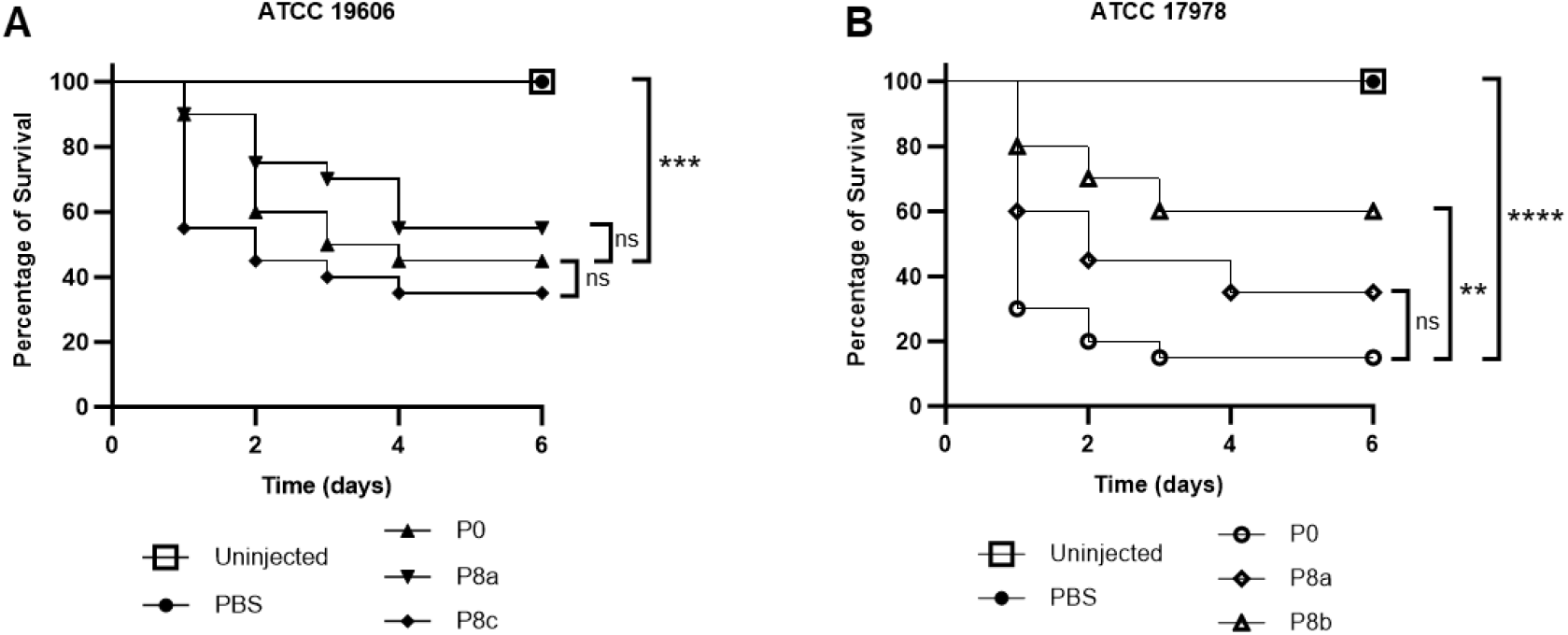
Development of resistance does not strongly affect *A. baumannii* virulence. Virulence of resistant isolates was compared to parental strains in a *Galleria mellonella* model. 10^6^ bacteria of the indicated strains were resuspended in 20 µl of PBS and were injected in a posterior proleg of 20 larvae for each strain. Injection of PBS alone was used as a control. Survival was assessed at the indicated time points. This was performed in ATCC19606 **(A)** and ATCC17978 **(B)** backgrounds in triplicate. Representative results are shown. GraphPad Prism was used to test for significant differences between the curves using the Log-rank (Mantel-Cox) test. Significance levels are indicated by asterisks (** = p-value < 0.01, *** = p-value < 0.001, **** = p-value < 0.0001).

## Discussion

AMPs have been identified as promising antimicrobial agents towards multidrug resistant (MDR) bacteria. Despite the recent progress made towards their clinical application, little is known on how pathogenic bacteria could adapt to a broad usage of AMPs. In this study, we investigated how one of the most successful nosocomial pathogens, *Acinetobacter baumannii*, adapts to in vitro treatment with the AMP TAT-RasGAP_317-326_. *A. baumannii* is known to rapidly evolve resistance and indeed clinical isolates are commonly MDR. In this study, we showed that *A. baumannii* is capable of acquiring resistance to TAT-RasGAP_317-326_. Importantly, in approximately half of the cases, we could observe emergence of cross-resistance to polymyxins. Since polymyxins are used as last resort antibiotics towards infections caused by MDR bacteria, appearance of such cross-resistance is highly problematic. In addition, we did not observe any significant reduction in bacterial fitness and virulence linked to emergence of cross-resistance. This indicates that the fitness cost induced by cross-resistance emergence is negligeable.

In a large majority of cross-resistant isolates, mutations in *pmrAB* genes were detected. PmrAB is a two-component system that was shown to regulate modifications of the LPS. Mutations affecting the activity of PmrAB were detected in polymyxins-resistant clinical isolates. Even though resistance selection towards TAT-RasGAP_317-326_ was performed in vitro, some of the mutations we obtained were identical to mutations detected in polymyxins-resistant clinical isolates (Fig. 4B, in bold). Selection of such mutations indicates that LPS may mediate affinity of both TAT-RasGAP_317-326_ and polymyxins with *A. baumannii*. This influence is however not affecting efficiency of all AMPs since mutants cross-resistant to TAT-RasGAP_317-326_ and polymyxins are not resistant to melittin. More experiments need to be done now with a broader set of AMPs and will allow to differentiate their modes of interaction with the bacterial surface.

Another interesting observation is the emergence of cross-resistance in approximately 50% of the isolates. This indicates that several mechanisms of resistance towards TAT-RasGAP_317-326_ can be selected, and only a subset of those results in cross-resistance to polymyxins. However, we observed no clear difference between cross-resistance and specific resistance regarding their influence on fitness and virulence. This would explain why both can occur at a similar rate.

In a few cases, cross-resistance was observed in absence of mutations in *pmrAB* genes. In these cases, whole genome sequencing allowed us to identify mutations possibly involved in cross-resistance development. Most promising candidates are a point mutation in *lptC*, encoding a LPS transporter subunit, and a mutation of the start codon of *crp*, encoding a cyclic AMP receptor protein. Further investigations are needed to determine the exact role of these mutations in the development of cross-resistance.

Despite several attempts, we never obtained cross-resistance in another Gram-negative bacterium, *Escherichia coli*. Previous attempts were performed using the MG1655 strain, whose LPS does not contain O-antigen. This incomplete LPS might hinder the development of cross-resistance. Alternatively, it could be that genome plasticity of *E. coli* is lower, making mutations less frequent. This may cause lower diversity of the outcomes during resistance selection. This needs to be investigated, possibly through artificially increasing the mutational rate of *E. coli* via mutagenic agents, for example. *E. coli* strains synthesizing a complete LPS containing the O-antigen could also be studied in order to determine what factors make cross-resistance development less favourable in *E. coli*.

Taken together, our results show the strong adaptability of bacteria towards antimicrobial agents. This adaptability can lead to development of cross-resistance that can be highly deleterious for the clinical usage of some antimicrobial agents. This needs to be considered before a new antimicrobial agent is released and used broadly in clinical settings. It will allow to determine the best strategy of treatment, such as combinations with other drugs and/or appropriate dosage, while also avoiding the emergence of unwanted resistance.

## Material and methods

### Bacterial strains, culture conditions and antimicrobial agents

All strains used in this study are listed in Table S3. Bacteria were grown overnight at 37°C with 200 rpm shaking in LB medium (10 g/L tryptone, 5 g/L yeast extract and 10 g/L NaCl), or statically on LB agar plate (LB supplemented with 15 g/L agar). TAT-RasGAP_317-326_ retro-inversed antimicrobial peptide (amino acid sequence DTRLNTVWMWGGRRRQRRKKRG) was provided by SBS Genetech (Beijing, China). Polymyxin B, apramycin, and spectinomycin were from Sigma-Aldrich brand (Merck, Darmstadt, Germany), gentamicin, tetracycline, and kanamycin by Applichem GmbH (Darmstadt, Germany), melittin by Enzo Life Science (Farmingdale, NY, USA) and colistin by LGC (Teddington, UK).

### Minimal inhibitory concentration (MIC) measurements

Overnight cultures were diluted to OD_600_=0.1 in fresh LB and incubated 1 hour at 37°C with shaking. Cultures were then diluted 10x and 10 µl of it was exposed to 2-fold serial dilutions of antimicrobials in LB in 96-well plates (in a total of 100 µl per well). After an incubation of approximately 18h at 37°C in static conditions and humidified atmosphere, OD_600_ was measured using a FLUOstar Omega microplate reader (BMG Labtech, Ortenberg, Germany). The MIC was defined as the lowest concentration of the antimicrobial causing at least 90% decrease of bacterial density.

### In vitro resistance selection

Overnight cultures were diluted 100x in 500 µl of LB containing 0.5 MIC of the indicated antimicrobial agent and incubated under shaking at 37°C overnight. Similar dilution was then performed in medium containing the same concentration of antimicrobial agent or a higher concentration. Culture in which bacterial growth could be detected was further diluted similarly as before. This was repeated for a total of 8 passages.

### Growth curves

Overnight cultures were diluted in fresh LB supplemented or not with 0.4% glucose or 2% NaCl to obtain an OD_600_=0.01. 250 µl of the bacterial suspension was transferred in a 96-well polypropylene plate. The OD_600_ was measured every 30 min during 16 hours at 37°C with shaking using a BioTek, EPOCH 2 microplate spectrophotometer (Agilent, Santa Clara, CA, USA).

### Microscopy

Overnight cultures were diluted to an OD_600_ of 0.1 and incubated for two hours at 37°C. Cultures were then centrifuged at 5000 rpm for 2 min, supernatant was discarded, and bacterial pellet was resuspended in the remaining medium. 5 µl of bacterial suspension was then mounted on a microscopy slide. Pictures were taken using a light AXIO Imager 2 microscope (Zeiss, Oberkochen, Germany) using an 100x oil immersion objective and the ZEN software. Images were then treated using ImageJ software (29).

### Whole genome sequencing and Sanger sequencing

Genomic DNA was extracted from overnight cultures using Wizard genomic purification kit (Promega, Madison, WI) and quantified using the Qubit system (Thermo Fisher Scientific, Waltham, MA, USA). A Nextera XT Kit (Illumina, San Diego, CA) was then used to produce libraries, which quality was controlled by Fragment Analyzer AATI (Agilent). Sequencing was performed on a MiSeq system using MiSeq Reagent Kits v2 (Illumina). Reads were assembled with spades v. 3.11.1 (30) and mapped on the corresponding reference genomes with bwa v. 0.7.17 (31). Variant calling was performed using gatk 4.0.2.0 (32). Identified SNPs were manually checked by visualizing the mapping with JBrowse (33).

Verification of mutations in specific genes was performed by amplification of the gene of interest by colony PCR. Primers upstream and downstream of the region of interest (Listed in Table S3) were designed using Geneious (Biomatters, Auckland, New Zealand) and PCR mix was prepared by combining 12.5 µl of 2x Taq MasterMix Dye (Promega) and 1 µl of 10 µM dilutions of each primer, in a total volume of 25 µl. A single colony of the strain of interest was resuspended in the mix and PCR protocol in a Biometra TRIO Thermocycler (Labgene Scientific, Châtel-Saint-Denis, Switzerland) using the following parameters: 95°C 5 min; 40 cycles of 95°C 30 s, 54°C 30 s, 72°C (1 min for every kbp of product); and finally, 72°C for 10 min. Proper amplification and correct size of PCR products were checked on a 1% agarose gel (120 V, 400 amp, 20 min). PCR products were purified using a QIAquick PCR purification system (Qiagen, Hilden, Germany) and sent to Microsynth (Balgach, Switzerland) for Sanger sequencing. Resulting sequences were aligned to the reference gene using Geneious software.

### Biofilm formation assays

Biofilm formation assays were mostly performed as described earlier (18). Overnight bacterial cultures were diluted 1:50 into fresh LB and further incubated for 4 hours. Bacteria were then washed two times with PBS pH 7.4 and resuspended in freshly prepared BM2 medium (62 mM potassium phosphate buffer, 7 mM ammonium sulfate, 10 µM iron sulfate, 40% glucose, 50% casein peptone, 2 mM magnesium sulfate) to obtain a final OD_600_ of 0.1. 100 µl of bacterial suspension was then transferred into a 96-well polypropylene plate and 250 in 24 well-plates containing glass coverslips. The plate was incubated 24 hours at room temperature in humidified atmosphere. 24-well plates were tilted with a 45° angle during the whole incubation time.For biomass quantification, the 96-well polypropylene plate was washed two times with nanopure H_2_0 and 125 µl of 0.1% crystal violet (Sigma, V5265) was added into each well. After 10 min incubation, the plate was washed again three times with water, and dried overnight. Accumulated dye was then resuspended with 30% acetic acid for 10 minutes and transferred into a new 96-well plate. Absorbance at 600 nm was then measured using a FLUOstar Omega microplate reader. The 24-well polypropylene plates were washed two times with PBS and labelled with LIVE/DEAD BacLight Kit (Invitrogen) following the manufacturer’s instructions for 15 min in the dark. Following two washes with PBS, bacteria were fixed with 4% paraformaldehyde, washed two more times with PBS and mounted on a glass slide using Moewiol resin. Biofilms were imaged using the Z-stacks option of a confocal microscope (LSM900, Zeiss) with a 40x oil immersion objective. Orthogonal projections were reconstructed using the ZEN software version 3.0.

### CRISPR-Cas9-based targeted mutagenesis

Point mutations were inserted in the genome of *A. baumannii* strain ATCC 17978 using the CRISPR-Cas9 genome editing method described by Wang et al. (22) with some adaptations and optimizations. All oligonucleotides used in this experiment were designed using Geneious software, synthesized by Microsynth AG and are listed in Table S3. Forward and reverse oligonucleotides encoding the desired gRNA (1 µl of Spacer-F and Spacer-R) were phosphorylated with a T4 polynucleotide kinase (New England BioLabs, Ipswich, MA, USA) in a total volume of 50 µl for 1 hour at 37°C. 0.5 µl of NaCl 5 M was added to the phosphorylated product and annealing was performed by incubating the sample at 95°C for 5 min, followed by a gradual decrease in temperature at a rate of 1°C per 10 seconds until reaching 25°C. Cloning of the gRNA encoding sequence in the pSGAb-km expression plasmid was performed using Goldengate assembly with 1 µl of 20-fold diluted annealed oligonucleotides, 50 ng of plasmid, 0.5 µl of T4 DNA ligase and 0.5 µl of Bsal HFv2 (New England Biolabs) in a total volume of 10 µl. Goldengate assembly was performed using a thermocycler with the following program: 25 cycles (3 minutes at 37°C and 4 minutes at 16°C) followed by 5 minutes at 50°C and 10 minutes at 80°C. 5 µl of the product was then transformed into thermocompetent DH5α and colonies were screened by PCR using the M13R and Spacer-F primers. Plasmid containing the correct insert was then harvested by miniprep (GeneJET Plasmid Miniprep Kit, Thermo Fisher Scientific).An overnight culture of *A. baumannii* ATCC 17978 strain baring the pCasAb-apr plasmid was diluted 100x into 50 mL of LB with apramycin and incubated at 37°C until OD_600_ reached 0.1-0.15. Cas9 expression was induced by addition of 1 mM IPTG (Applichem), and culture was further incubated for two hours. Bacteria were then washed three times with ice cold milliQ water and electroporation was performed with 100 ng or 200 ng of the pSGAb-km plasmid and 3 µl or 6 µl of the donor ssDNA (100 µM) using chilled 1 mm electroporation cuvette and a Gene Pulser Xcell Electroporation system (BioRad, Hercules, CA, USA). After electroporation, cells were immediately resuspended into 1 mL of ice-cold LB and incubated 1 hour at 37°C with shaking. Transformed cells were plated on LB agar plates supplemented with the corresponding antibiotics (apramycin and kanamycin) and incubated overnight. The gene of interest was amplified by colony PCR. The PCR product was purified with a QIAquick PCR purification kit (Qiagen) and sent for Sanger sequencing (Microsynth).

### *Galleria mellonella* infection model

Overnight cultures of the indicated *A. baumannii* strains were washed with PBS and serially diluted and plated on LB agar petri dishes. The number of living bacteria per ml was measured by Colony Forming Units quantification. OD_600_ of the culture was measured in parallel and CFU/OD_600_ was then calculated. Bacteria were diluted to obtain a final suspension of bacteria corresponding to 10^6^ bacteria in 20 μl of PBS. 20 μl of the suspension were then injected in a posterior proleg of *Galleria Mellonella* (Bait Express, Basel, Switzerland). 20 larvae were injected per group, with control groups being either uninjected or injected with PBS. Larvae were then incubated at 37°C and survival was monitored daily up to 6 days. Survival graphs and statistical analyses (Mantel-Cox test) were performed using GraphPad Prism software version 10.1.2 (La Jolla, CA).

## Acknowledgments

We would like to thank Sébastien Aeby for technical support and Gilbert Greub for providing lab space and sharing equipment. Clinical strains were provided by the bacteriology unit of Institute of Microbiology. Participation of Master students to this work was financed by the School of Biology, Faculty of Biology and Medicine of the University of Lausanne. Sequencing analyses were performed by the group of Claire Bertelli at the Institute of Microbiology and financed by a personal grant from F. Paulsen. We especially thank Maria Georgieva for her careful proofreading of this article.

