## Supplementary figures for "In vitro resistance selection in *Acinetobacter baumannii* against the antimicrobial peptide TAT-RasGAP_317-326_ can cause cross-resistance to polymyxins through mutations in *pmrAB*"

**Figure S1. Resistance acquisition does not induce in vitro growth defects in presence of glucose.** P0 and P8 isolates upon selection with TAT-RasGAP$_{317-326}$ **(A)**, polymyxin B **(B)** or tetracycline **(C)** were grown overnight in LB and diluted to an $OD_{600}$ of 0.01 in fresh LB supplemented with 0.4% glucose. Growth was then assessed by $OD_{600}$ measurement each 30 minutes for a total of 16 hours. Values are the average of three experiments. Error bars represent standard deviation of the three replicates.

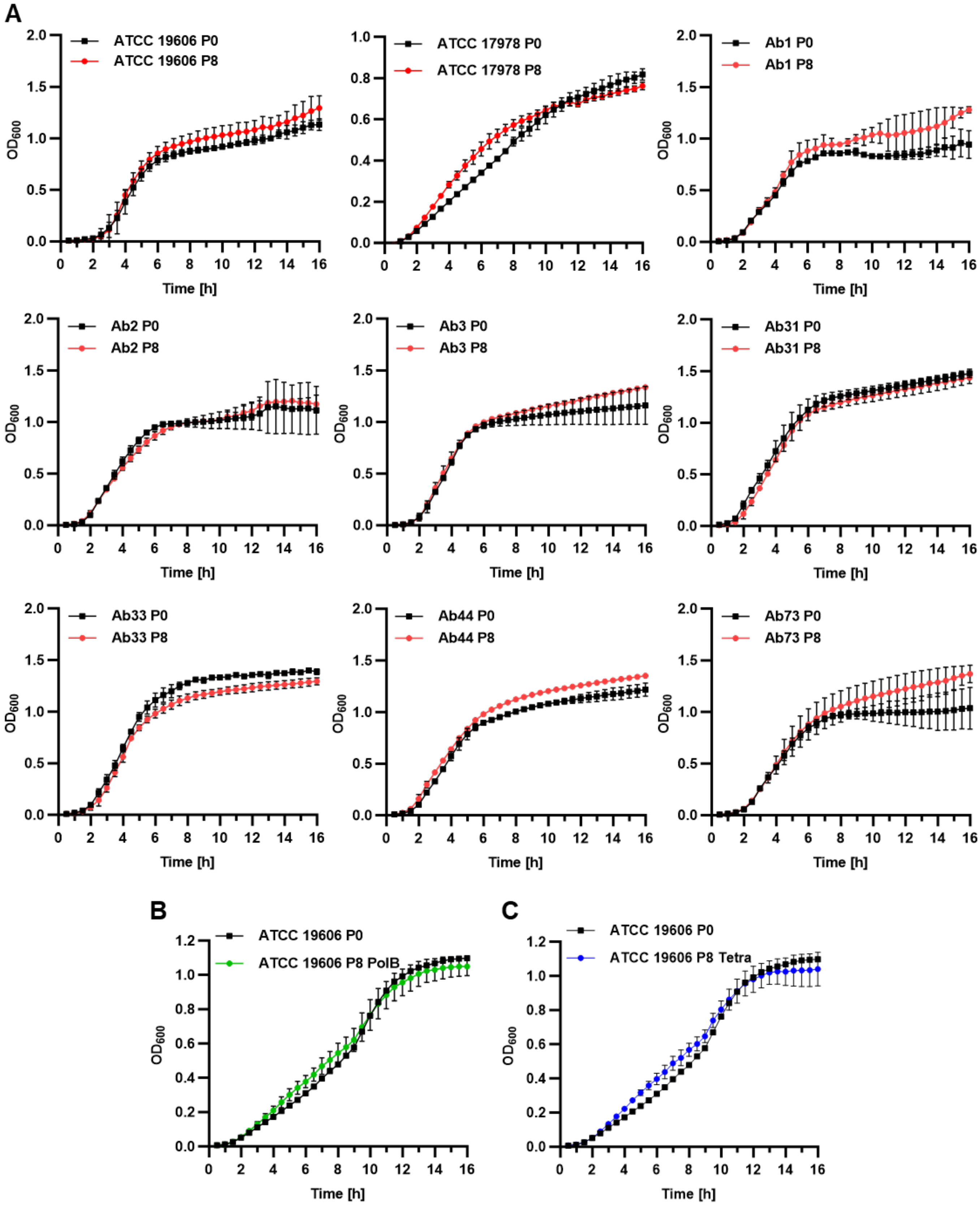

**Figure S2. Resistance acquisition does not induce in vitro growth defects in presence of increased salt concentration.** P0 and P8 isolates upon selection with TAT-RasGAP$_{317-326}$ **(A)**, polymyxin B **(B)** or tetracycline **(C)** were grown overnight in LB and diluted to an $OD_{600}$ of 0.01 in fresh LB containing 2% of NaCl. Growth was then assessed by $OD_{600}$ measurement each 30 minutes for a total of 16 hours. Values are the average of three experiments. Error bars represent standard deviation of the three replicates.

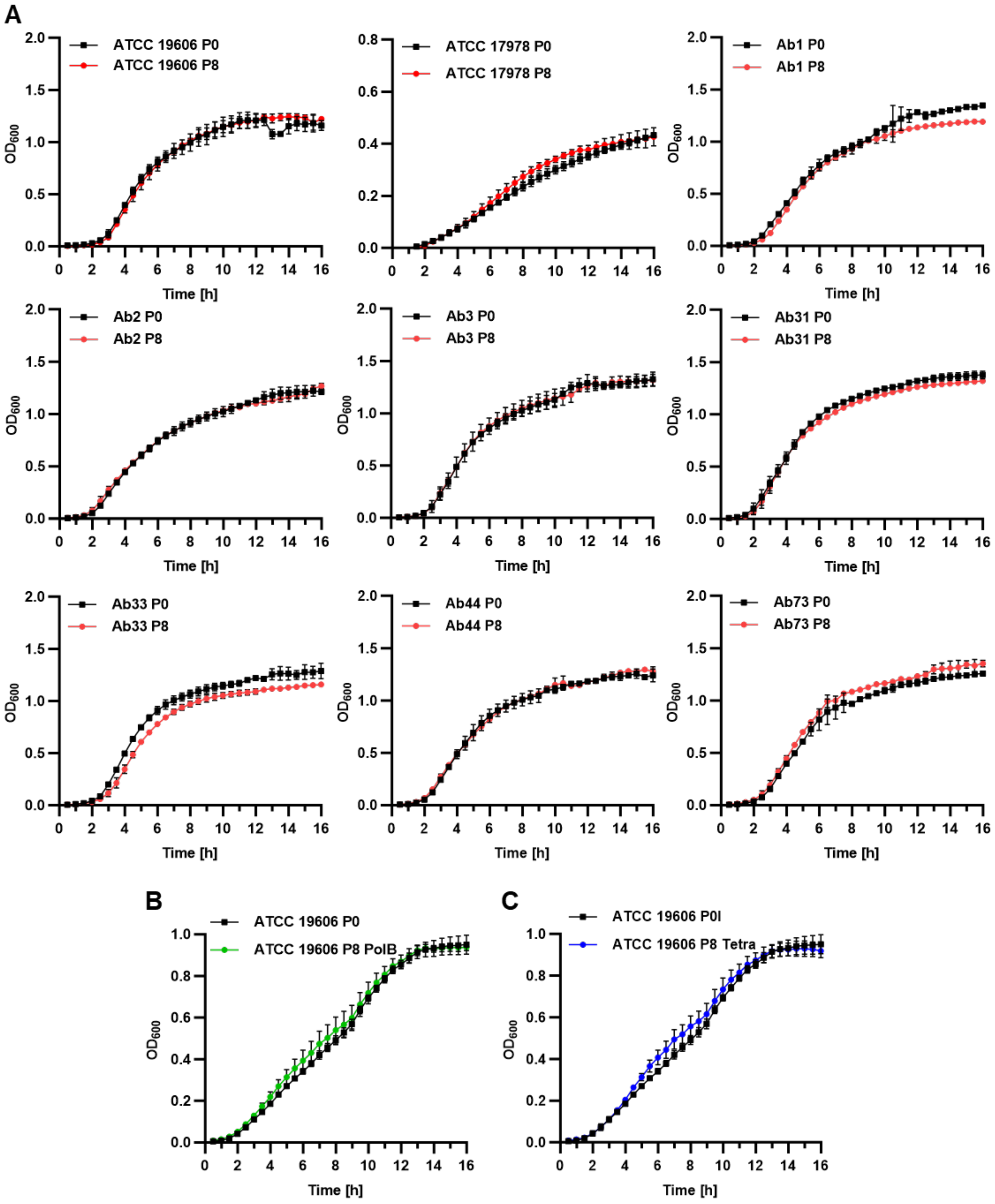

**Figure S3. Resistance acquisition does not induce morphology in vitro.** P0 and P8 isolates upon selection with TAT-RasGAP$_{317\text{-}326}$, polymyxin B or tetracycline were grown overnight in LB, diluted to an $OD_{600}$ of 0.1 in fresh medium and grown for 2 hours. Morphology of the different isolates was then assessed by microscopy.

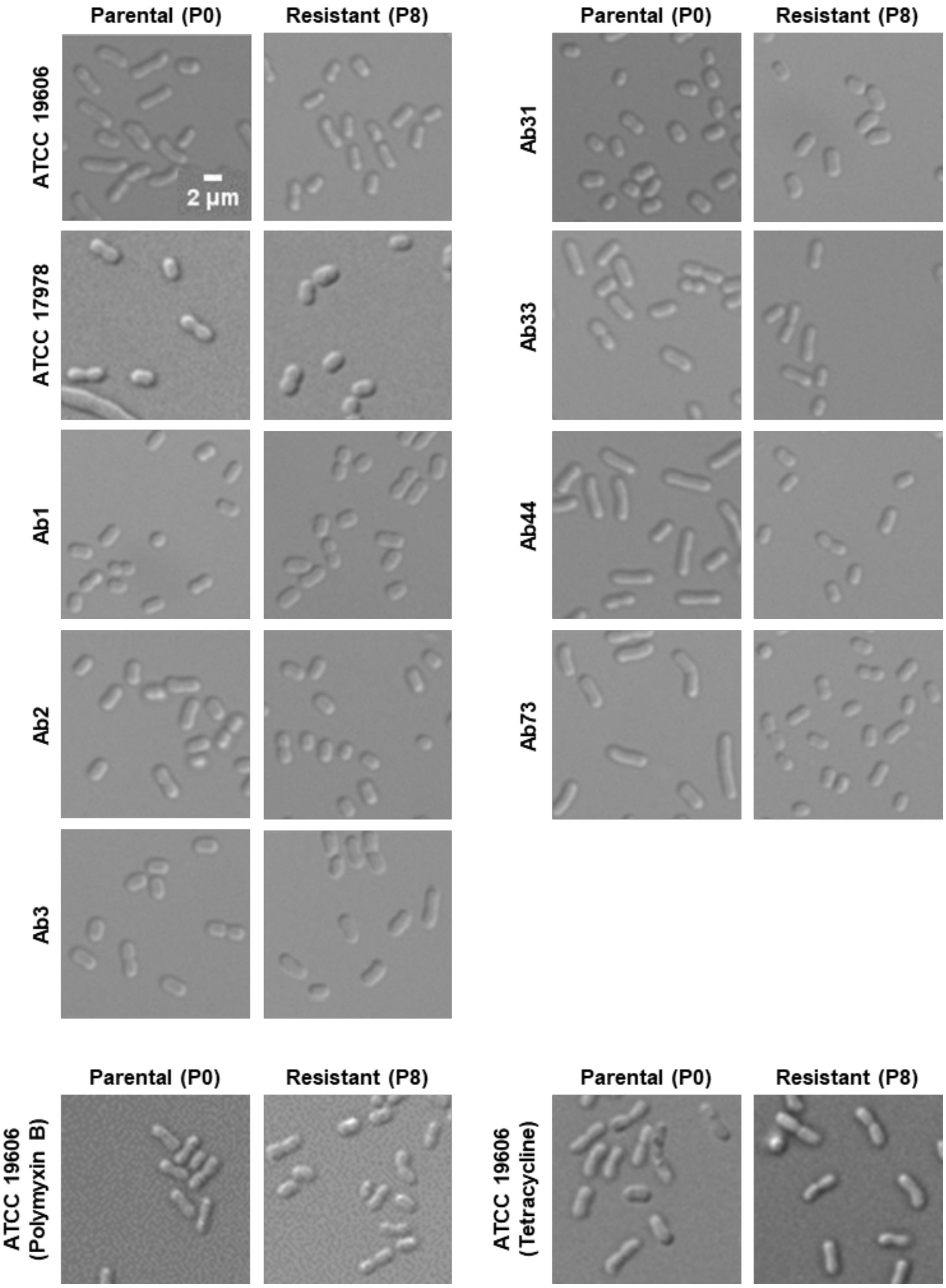

**Figure S4.** The effect of resistance acquisition on the structure of the biofilm was assessed by live-dead staining followed by Z-stacks acquisition with a confocal microscope. Reconstruction of Z projections are shown on the top and on the right for each strain. This figure is a complement to Figure 3.

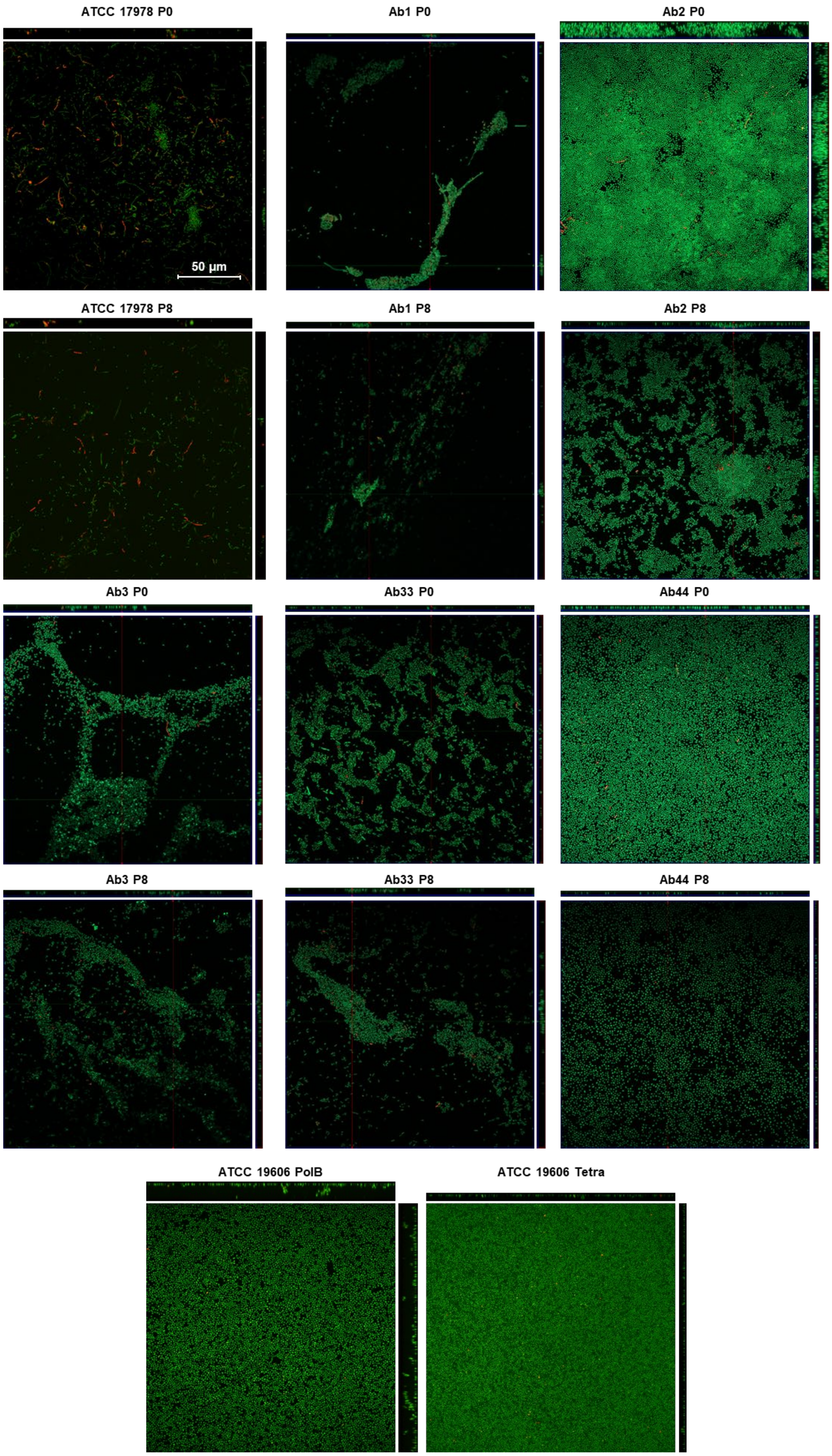

**Figure S5. MIC of TAT-RasGAP$_{317-326}$ (Black) and polymyxin B (Red) of all resistant strains selected in this study.** MICs were measured on the indicated strains in duplicates. Black lines represent the mean of the replicates.

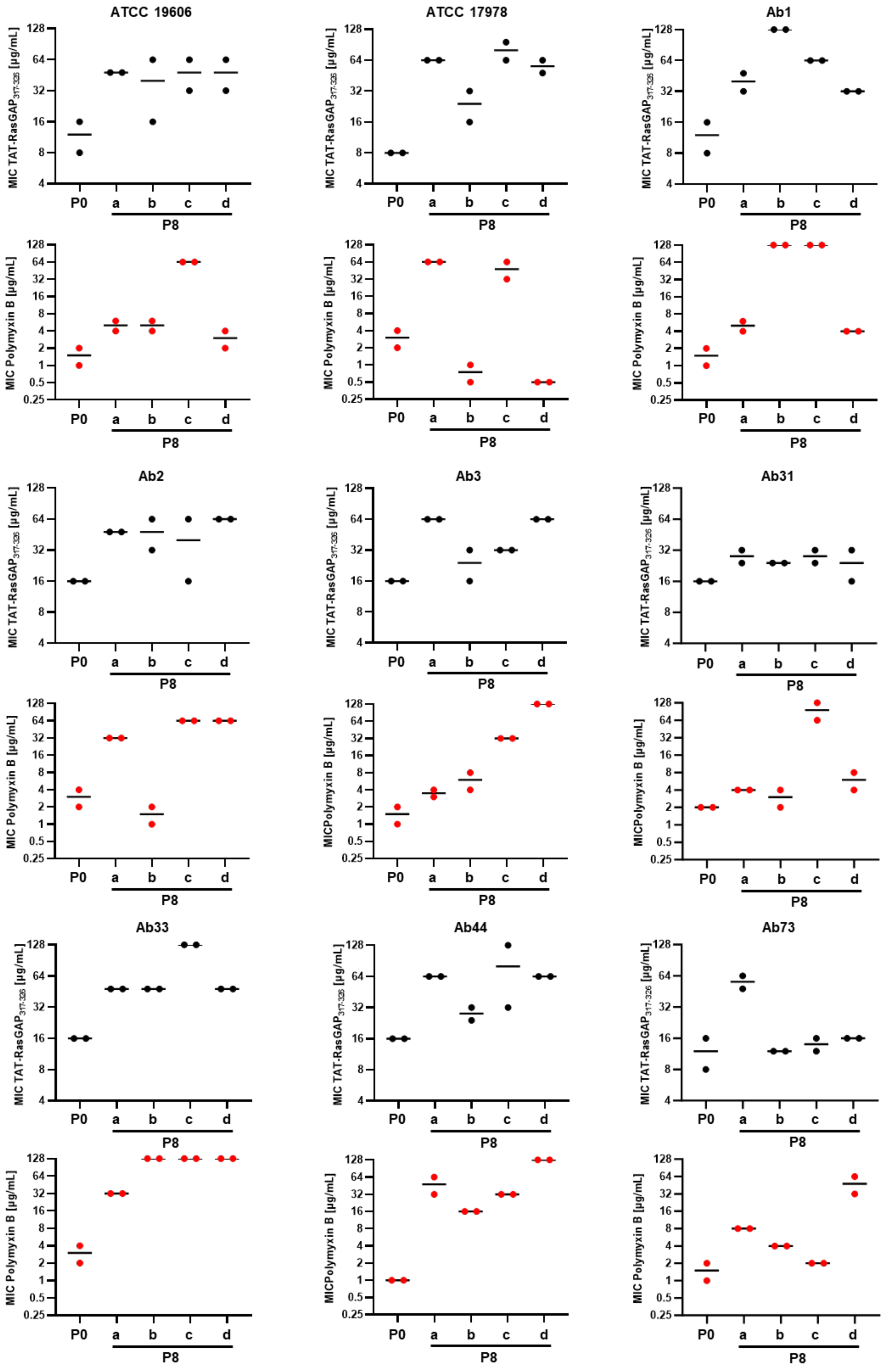

**Figure S6. Cross-resistance to polymyxin B is stable upon 8 passages in absence of drug selection.** ATCC19606 parental strain and selected resistant isolates b, c and d were passaged in absence of any antimicrobial agents for a total of 8 passages. MICs of TAT-RasGAP$_{317\text{-}326}$ and polymyxin B before (Resistant) and after (Back-selection) the 8 passages were measured.

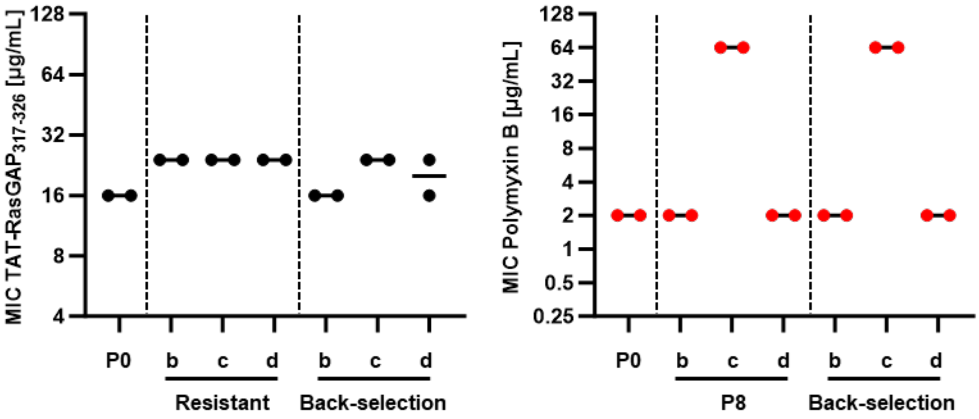
